# Intracellular Oxidative Stress Levels are Significantly Associated with the Green Autofluorescence Intensity of Buthionine Sulfoximine-Treated B16-F10 Cells

**DOI:** 10.1101/2021.03.02.433583

**Authors:** Wanzhi Tang, Weihai Ying

## Abstract

Since oxidative stress is a critical common pathological factor of numerous diseases, it is critical to find biomarkers for non-invasive evaluations of the levels of oxidative stress in the body. Our previous studies have indicated that epidermal green autofluorescence (AF) is a novel biomarker of this type: The oxidative stress inducer buthionine sulfoximine (BSO) can dose-dependently increase the epidermal green AF of mice, with BSO doses being significantly associated with the AF intensity. However, it is necessary to use skin cell cultures to investigate the mechanisms underlying the relationships between BSO and the green AF intensity. In our current study we found that BSO concentration-dependently increased the green AF intensity of B16-F10 cells a skin cell line, with BSO concentrations being significantly associated with the AF intensity. BSO also concentration-dependently increased the intracellular DCF signals an index of ROS levels. The green AF intensity of the cells was also significantly associated with the intracellular ROS levels. Moreover, we found that the green AF intensity was significantly associated with the cell death induced by BSO. Collectively, our study has provided first evidence indicating that the green AF intensity of skin cells is significantly associated with both intracellular ROS levels and cell death of the skin cells exposed to oxidative stress, which has indicated that green AF is a novel biomarker for both oxidative stress and cell death.

## Introduction

Oxidative stress is a critical common pathological factor of numerous diseases, including acute ischemic stroke (AIS) (1, 2), cardiovascular diseases (3), lung cancer (4), Alzheimer’s disease (5) and Parkinson’s disease (6). Current approaches for determining the levels of oxidative stress in the body require blood drawing, which are invasive and relatively time consuming (7). It is critical to find biomarkers for non-invasive evaluations of the levels of oxidative stress in the body. Moreover, increased oxidative stress is a general index for the pathological state of a person (8, 9). For regular monitoring and management of a person’s health state, it is also crucial to discover biomarkers and approaches for non-invasive evaluations of the levels of oxidative stress in the body.

Human autofluorescence (AF) from the advanced glycation end-product (AGE)-modified collagen and elastin has shown promise for non-invasive diagnosis of diabetes and diabetes-related pathology (10). Our previous studies have indicated that epidermal green autofluorescence (AF) is a novel biomarker for non-invasive evaluations of the levels of oxidative stress in the body: BSO is an inhibitor of γ-glutamylcysteine synthetase - an important enzyme for GSH synthesis, which can lead to increased oxidative stress by inhibiting GSH synthesis (11). We have found that BSO can dose-dependently increase the epidermal green AF of mice, with BSO doses being significantly associated with the AF intensity (12). Our study has also indicated that increased skin’s green AF is a promising biomarker for diagnosis of such major diseases as AIS (13), lung cancer (14) and Parkinson’s disease (15). However, regarding our finding that BSO induced increased epidermal green AF of mice, the following important scientific questions have remained unanswered: 1) Does BSO induce increased green AF of skin cells by its direct effect on the skin cells? 2) Are intracellular oxidative stress levels of skin cells associated with the green AF intensity of the skin cells? 3) Is the oxidative stress-induced green AF intensity of the skin cells associated with the levels of the skin cell death ?

In order to obtain answers to these questions, it is necessary to conduct skin cell cultures studies. Therefore, in our current study we determined the relationships among BSO concentrations, green AF intensity of the skin cells, intracellular oxidative stress levels of the skin cells, and the cell death by using B16-F10 cells as a skin cell model. We found significant associations among these important factors.

## Materials and Methods

### Materials

All chemicals were purchased from Sigma-Ardrich (St. Louis, MO, USA) unless otherwise expressly noted.

### Cell Cultures

B16-F10 cells were plated into 24-well or 6-well culture plates in Dulbecco’s modified Eagle medium (HyClone, Logan, UT, United States) supplemented with 10% fetal bovine serum (Gibco, Carlsbad, CA, United States), 100 units/ml penicillin and 100 μg/ml streptomycin at 37 °C in a humidified incubator under 5% CO_2_.

### Imaging of the green autofluorescence of B16-F10 cells

The AF signals of the cells were observed under a Leica confocal fluorescence microscope with an excitation wavelength of 488 nm and an emission wavelength of 500-530 nm.

### Determinations of ROS

For Dichlorofluorescin (DCF) assay, 2,7-Dichlorofluorescin diacetate (DCFH-DA, Beyotime, China), a reactive oxygen species (ROS)-specific fluorescent probe (16), was used to measure total intracellular ROS levels. Cells were incubated with 20 μM DCFH-DA dissolved in DMEM without fetal bovine serum (FBS) for 20 min at 37°C in a dark room. After 3X washes with PBS, the cells were analyzed by flow cytometry (FACSAria; Becton Dickinson, Heidelberg, Germany) to detect the mean fluorescence intensity (MFI) with an excitation wavelength of 488 nm and an emission wavelength of 525 nm.

### Determinations of GSH and GSSG levels

The concentrations of total glutathione, GSH and GSSG of cells were determined according to the protocol of a GSH / GSSG Assay Kit (Beyotime, China).

### Extracellular lactate dehydrogenase (LDH) and Intracellular LDH assays

Cell survival was quantified by measuring the intracellular LDH activity of the cells, while the extracellular LDH activity was used to quantify cell death. Extracellular LDH activity and intracellular LDH activity were determined by determining the LDH activity of the cell culture medium and the cell lysates, respectively. Cell lysates were prepared by the following procedures: Cells were lysed for 15 min in lysing buffer containing 0.04% Triton X-100, 2 mM HEPES and 0.01% bovine serum albumin (pH 7.5). For LDH activity assays, 50 μL cell culture medium or 50 μL cell lysates was mixed with 150 μL 500 mM potassium phosphate buffer (pH 7.5) containing 0.34 mM NADH and 2.5 mM sodium pyruvate. The A340nm changes were monitored over 90 sec. Percentage of cell survival was calculated by normalizing the LDH values of samples to LDH activity measured in the lysates of control (wash only) culture wells.

### Statistical analyses

All data are presented as mean ± SEM. Data were assessed by one-way ANOVA, followed by Tukey post hoc test. *P* values less than 0.05 were considered statistically significant.

## Results

We determined the green AF of B16-F10 cells treated with various concentrations of BSO: BSO induced dose-dependent increases in the green AF intensity, assessed 36 h after the BSO treatment (Figs. 1A and 1B). The BSO concentrations were also significantly associated with the green AF intensity (Fig. 1C).

**Fig. 1.**
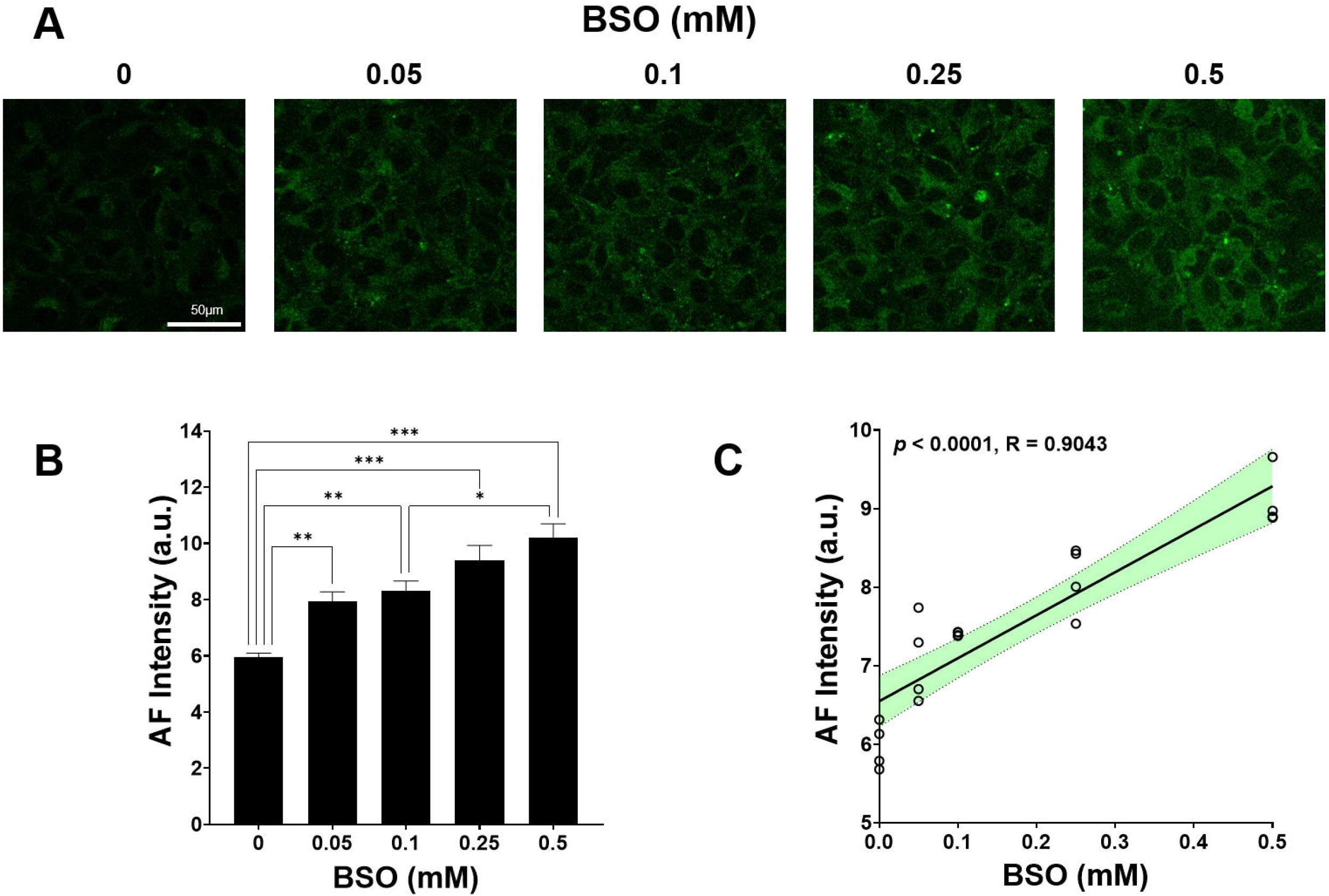
The BSO concentrations were significantly associated with the green AF intensity of BSO-treated B16-F10 cells. (A) Confocal images of the green AF of the B16-F10 cells treated with various concentrations of BSO, assessed 36 h after the BSO treatment. Scale bar = 50 μm. (B) Quantifications of the green AF intensity of the confocal images showed that BSO produced concentration-dependent increases in the green AF intensity. N = 8. *, *P* < 0.05; **, *P* < 0.01; ***, *P* < 0.005. (C) The BSO concentrations were significantly associated with the green AF intensity.

We also determined the intracellular (ROS) levels of the BSO-treated B16-F10 cells by using DCF fluorescence as a ROS index: BSO induced concentration-dependent increases in the DCF fluorescence signals, assessed 36 h after the BSO treatment (Figs. 2A and 2B). The BSO concentrations were significantly associated with the DCF fluorescence signals (Fig. 2C). The green AF intensity was also significantly associated with the DCF fluorescence signals (Fig. 2D). Similar results were also obtained by our flow cytometry assays using DCFH-DA as fluorescent probe (Figs. 2E - 2H). The effects of BSO on the levels of total glutathione, GSH and GSSG of the cells were also determined: BSO induced profound decreases in the levels of total glutathione, GSH and GSSG (Figs. 3A and 3B).

**Fig. 2.**
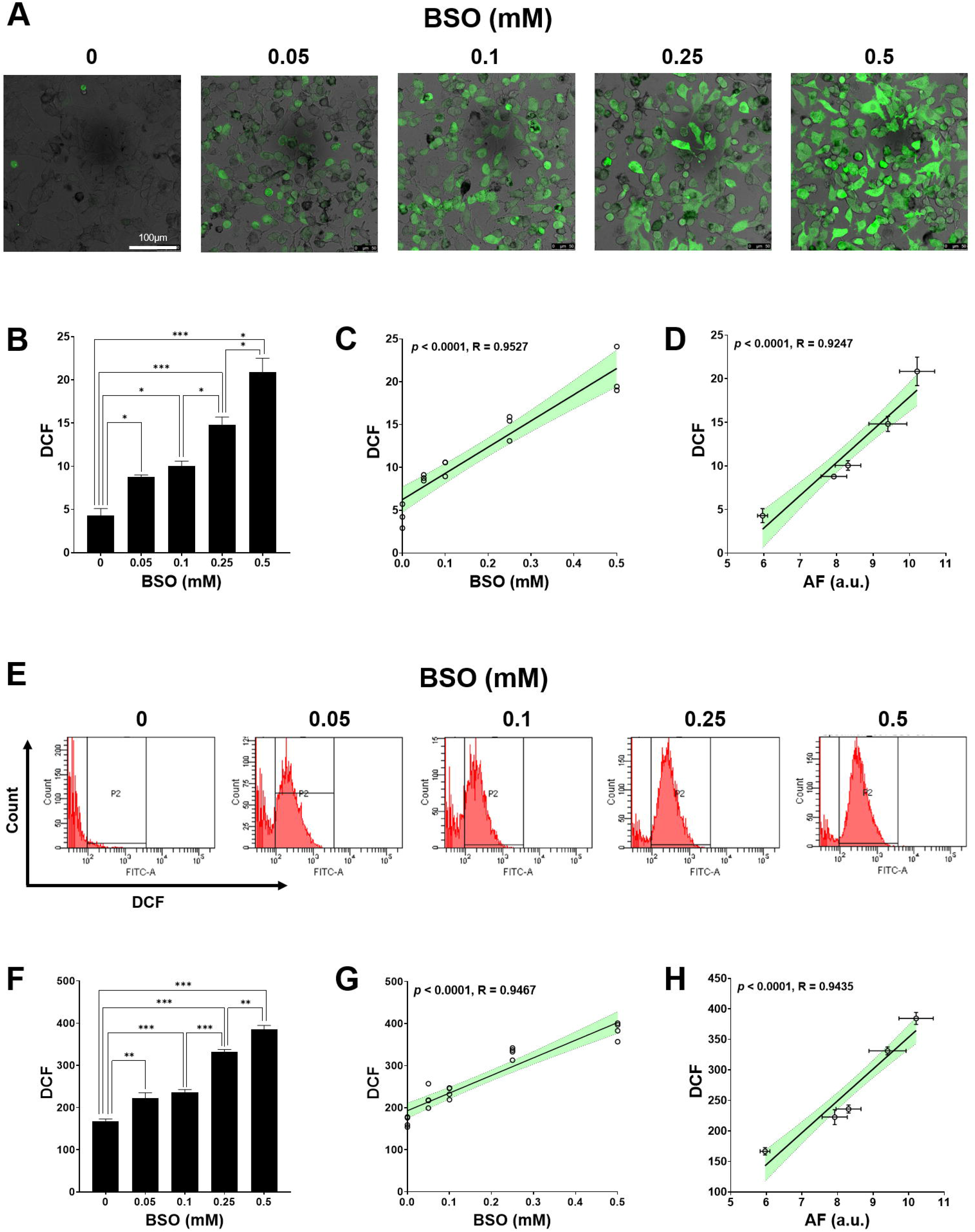
The green AF intensity was significantly associated with the intracellular ROS levels of BSO-treated B16-F10 cells. (A) The merged images of light-field images and fluorescence images showed the DCF fluorescence signals of BSO-treated B16-F10 cells. Scale bar = 100 μm. (B) Quantifications of the DCF fluorescence intensity of the fluorescence images showed that BSO produced concentration-dependent increases in the DCF fluorescence intensity. N = 6. *, *P* < 0.05; **, *P* < 0.01; ***, *P* < 0.005. (C) The BSO concentrations were significantly associated with the DCF fluorescence intensity of the BSO-treated cells. (D) The DCF fluorescence intensity was significantly associated with the green AF intensity of the BSO-treated cells. (E) The flow cytometry assays using DCFH-DA as a probe showed the DCF fluorescence signals of B16-F10 cells treated with various concentrations of BSO. (F) Quantifications of the DCF fluorescence intensity determined by the flow cytometry assays showed that BSO produced concentration-dependent increases in the DCF fluorescence intensity. N = 6. *, *P* < 0.05; **, *P* < 0.01; ***, *P* < 0.005. (G) BSO concentrations were significantly associated the DCF fluorescence intensity determined by the flow cytometry assays. (H) The DCF fluorescence intensity determined by the flow cytometry assays was significantly associated with the green AF intensity of the BSO-treated cells.

**Fig. 3.**
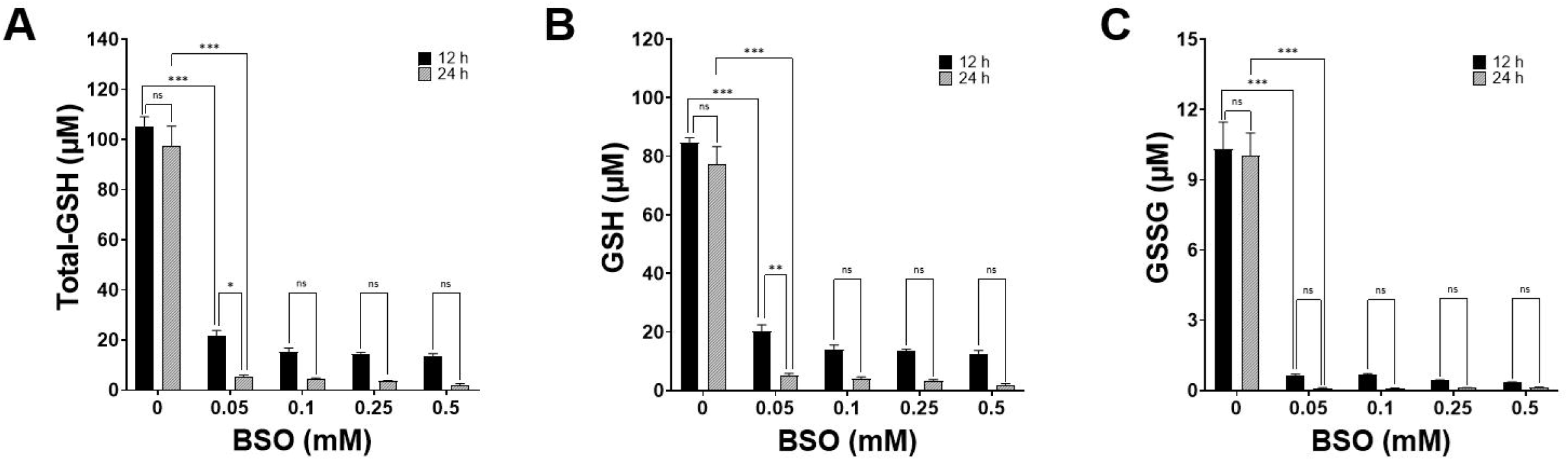
BSO induced profound decreases in the levels of total glutathione, GSH and GSSG. (A) BSO produced profound decreases in the intracellular levels of total glutathione of B16-F10 cells. (B) BSO produced profound decreases in the intracellular GSH levels of B16-F10 cells. (C) BSO produced profound decreases in the intracellular GSSG levels of B16-F10 cells. N = 6. *, *P* < 0.05; **, *P* < 0.01; ***, *P* < 0.005.

We further investigated the effects of BSO on the levels of both extracellular LDH activity and intracellular LDH activity of B16-F10 cells: BSO induced concentration-dependent increases in the extracellular LDH levels, assessed 36 h after the BSO treatment (Fig. 4A), with the green AF intensity being significantly associated with the extracellular LDH activity (Fig. 4B). BSO also induced concentration-dependent decreases in the intracellular LDH levels, assessed 36 h after the BSO treatment (Fig. 4C), with the green AF intensity being negatively associated with the intracellular LDH activity (Fig. 4D).

**Fig. 4.**
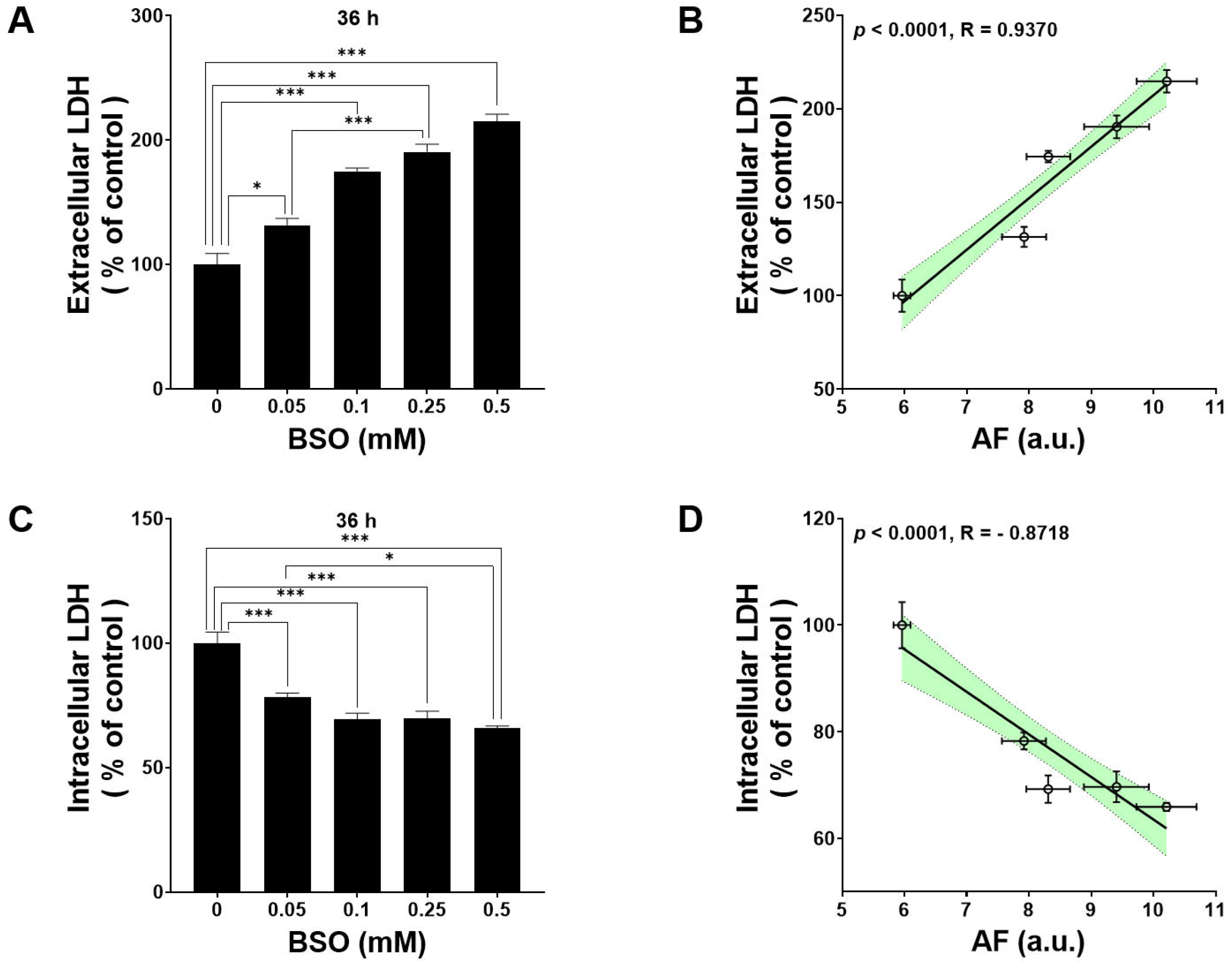
The green AF intensity was significantly associated with both extracellular LDH levels and the intracellular LDH levels of the BSO-treated B16-F10 cells. (A) BSO produced concentration-dependent increases in the extracellular LDH levels, assessed 36 h after the BSO treatment. (B) The green AF intensity was significantly associated with the extracellular LDH activity. (C) BSO produced concentration-dependent decreases in the intracellular LDH levels, assessed 36 h after the BSO treatment. (D) The green AF intensity was significantly associated with the intracellular LDH activity. N = 8. *, *P* < 0.05; **, *P* < 0.01; ***, *P* < 0.005.

### Discussion

Based on our previous finding that BSO can dose-dependently increase the epidermal green AF of mice, our current study has obtained the following important findings regarding the relationships among BSO concentrations, green AF intensity of the skin cells, intracellular oxidative stress levels of the skin cells, and the cell death by using B16-F10 cells as a skin cell model: First, BSO concentration-dependently increased the green AF intensity of B16-F10 cells, with BSO concentrations being significantly associated with the AF intensity; second, the green AF intensity of the cells was also significantly associated with the intracellular ROS levels of the cells; and third, the green AF intensity is significantly associated with the levels of cell death of BSO-treated skin cells.

Our current study has provided information for answering the important scientific questions related with our previous finding that BSO dose-dependently increased the epidermal green AF of mice: It has been unclear if BSO induced increased green AF of skin cells by its direct effect on the skin cells. Our current finding has indicated that BSO induced increased green AF of skin cells by its direct effect on increasing the intracellular oxidative stress of the skin cells, which has exposed an important mechanism underlying the BSO-induced increase in the epidermal green AF of mice. This finding has also indicated that the BSO-treated B16-F10 cells is a valuable cellular model for investigating the mechanisms underlying the BSO-induced increases in the epidermal green AF. Since we have found that the skin’s green AF intensity of the patients holds excellent promise for diagnosis of several oxidative stress-associated major diseases such as AIS (13), lung cancer (14) and Parkinson’s disease (15), findings regarding the mechanisms underlying the oxidative stress-induced increases in the skin’s green AF are of critical significance for establishing the novel AF-based diagnostic approaches of the diseases.

Our study has shown that the concentrations of extracellularly added BSO are significantly associated with the green AF of B16-F10 cells, while it is still important to determine if the intracellular oxidative stress levels in the BSO-treated cells are also significantly associated with the green AF intensity of the cells. Our study has shown that levels of DCF signals are significantly associated with the green AF intensity. This finding is also valuable for understanding the mechanism underlying the oxidative stress-induced increases in the epidermal green AF.

It has been unclear if the oxidative stress-induced green AF intensity of the skin cells is associated with the death of the skin cells. Our current finding has indicated that oxidative stress-induced green AF intensity of the skin cells is significantly associated with the death of the skin cells. This finding has indicated that the green AF intensity of the oxidative stress-exposed skin cells is not only a biomarker for oxidative stress, but also a biomarker for a major consequence of oxidative damage of cells - oxidative cell death. Based on this finding, new non-invasive approaches for evaluating the oxidative stress-induced skin cell death may be achieved.

## Acknowledgment

The authors would like to acknowledge the financial support by a Major Special Program Grant of Shanghai Municipality (Grant # 2017SHZDZX01) (to W.Y.).

